# Inhibition of p53 improves CRISPR/Cas - mediated precision genome editing

**DOI:** 10.1101/180943

**Authors:** Emma Haapaniemi, Sandeep Botla, Jenna Persson, Bernhard Schmierer, Jussi Taipale

**Author notes:** Corresponding authors: Jussi Taipale, Department of Medical Biochemistry and Biophysics, Karolinska Institutet, Scheeles väg 2, SE-171 77 Stockholm. Corresponding authors: Bernhard Schmierer, Department of Medical Biochemistry and Biophysics & Science for Life Laboratory, Karolinska Institutet, Scheeles väg 1, SE-171 77 Stockholm.

## Abstract

We report here that genome editing by CRISPR/Cas9 induces a p53-mediated DNA damage response and cell cycle arrest. Transient inhibition of p53 prevents this response, and increases the rate of homologous recombination more than five-fold. This provides a way to improve precision genome editing of normal cells, but warrants caution in using CRISPR for human therapies until the mechanism of the activation of p53 is elucidated.

## Main text

CRISPR/Cas9 has become a popular precision genome editing tool. It induces DNA double-stranded breaks (DSBs), which are repaired either by the error-prone non-homologous-end-joining (NHEJ), or precisely by the homologous recombination (HR) pathway. The choice of repair mechanism is dependent on the stage of the cell cycle, with NHEJ predominating in G1, and HR becoming efficient only during and after DNA replication. In precision genome editing, a repair template that is homologous to the cut locus is introduced to cells, where it is taken up by the endogenous HR machinery, leading to a scarless precision editing of the genome in an optimal case.

Precision genome editing by HR is very efficient in some tumor cell lines. By contrast, it has been very difficult to efficiently edit primary cells, which commonly undergo apoptosis, and/or preferentially employ NHEJ for damage repair, leading to a high ratio of indels to precise genome edits ^1^. Several efforts have been made to improve genome editing of primary cells, including increasing the concentration of the repair DNA template, delivering NHEJ inhibitors, and optimizing transfection ^1-5^. Although these means can improve precision editing up to threefold, a mechanistic explanation for its general inefficiency is lacking.

During the course of identifying essential genes in a large panel of cell lines using standard CRISPR/Cas9 “dropout” screens (Fig 1A; and Supplementary Methods)^6, 7^, we observed that guide-RNAs (gRNAs) targeting essential genes were not efficiently depleted in the human immortalized normal retinal pigment epithelial cell line RPE-1. Instead, we noted a dramatic increase of gRNAs targeting the tumor suppressor p53, its transcriptional target p21 (CDKN1A), and the retinoblastoma tumor suppressor gene (RB1) that is necessary for the cell cycle arrest induced by p21 (Fig 1B)^8^. The effect was not specific to RPE1 cells, as a similar, albeit weaker, effect was observed in another non-tumorigenic breast cell line MCF10A that has a partially active p53 pathway (not shown).

**Figure 1.**
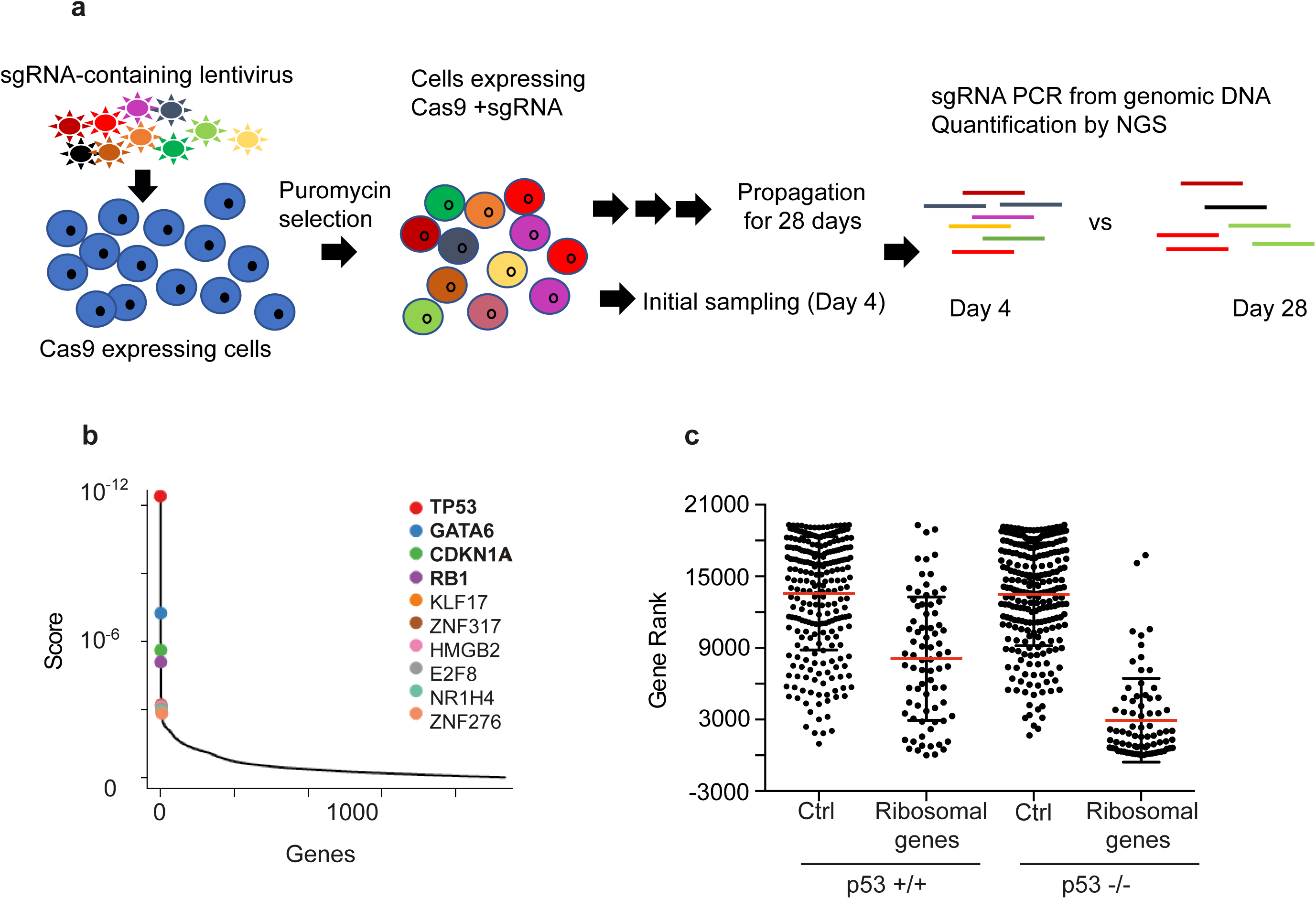
CRISPR screen is affected by p53 DNA damage response. **a)** Schematic representation of the screen setup. Cas9 expressing cells are transduced with a gRNA library in pool, propagated for several weeks, and finally sequenced. gRNAs targeting fitness genes are depleted during this time, while guides targeting growth inhibitory genes are enriched. **b)** gRNAs targeting p53 and its effectors p21 and RB1 were dramatically enriched (modified Robust Rank Aggregation analysis MaGeck^19^ in immortalized Retinal Pigment Epithelium (RPE1) cells. The enriched hits which are consistent between replicates are highlighted in bold. **c)** The gRNAs against ribosomal genes should deplete quickly from a pool, as their targets are essential for cell viability. Consequently, their gene rank is low. Contrary to this expectation, in p53^+/+^ cells the gRNAs targeting ribosomal genes are depleted randomly (left, median rank 8,034), whereas in p53^-/-^ cells, ribosomal gRNAs are among the first to drop out and concentrate in the lowest decile (right, median rank 1,627). Control gRNAs with no genomic targets behave similarly in p53 null and WT cells.

These observations suggested to us that in cells with a wild-type p53 response, the DSBs induced by Cas9 activate p53, leading to either growth arrest or apoptosis. This effect could also explain the failure to detect essential genes in the screen, as all cells containing a gRNA targeting a gene that is not on the p53 pathway are expected to arrest or die. To test this hypothesis, we carried out new genome-wide CRISPR/Cas9 screens in RPE-1 cells and in RPE-1 cells deficient in p53^9^ using a previously published Brunello library^10^. Consistently with the hypothesis, gene set enrichment analysis performed on the ranked list of genes^11, 12^ identified known essential pathways (general transcription and translation) in p53^-/-^ but not in p53^+/+^ cells (Fig 1C and Supplementary Table S1). Conversely, guides targeting p21 were enriched in p53^+/+^ but not in p53^-/-^ cells (4.7 fold versus 0.8 fold, respectively), indicating that loss of p21 abrogates the p53 dependent growth arrest, but has no proliferative effect of its own. We conclude that DSBs introduced by gene targeting using CRISPR/Cas9 trigger a p53 response, irrespective of the locus targeted, and that this effect severely hampers negative selection or dropout screening.

In lentiviral screens, the Cas9 and the gRNA are integrated into the genome and constitutively active. To assess whether a transient Cas9 activation would also trigger a p53 response, we transfected RPE-1 p53^+/+^ and p53^-/-^ cells with ribonucleoprotein (RNP) complexes containing a guide previously reported to have no off-targets^13^. We found that even such a short exposure to Cas9/gRNA activity resulted in a marked upregulation of p21 (Fig 2A) and a concomitant G1 arrest (Fig. 2B) in p53^+/+^, but not p53^-/-^, cells. This G1 arrest is likely transient, as we did not observe increased apoptosis as measured by Caspase 3 cleavage after 24 hours of RNP delivery (Fig. 2C).

**Figure 2.**
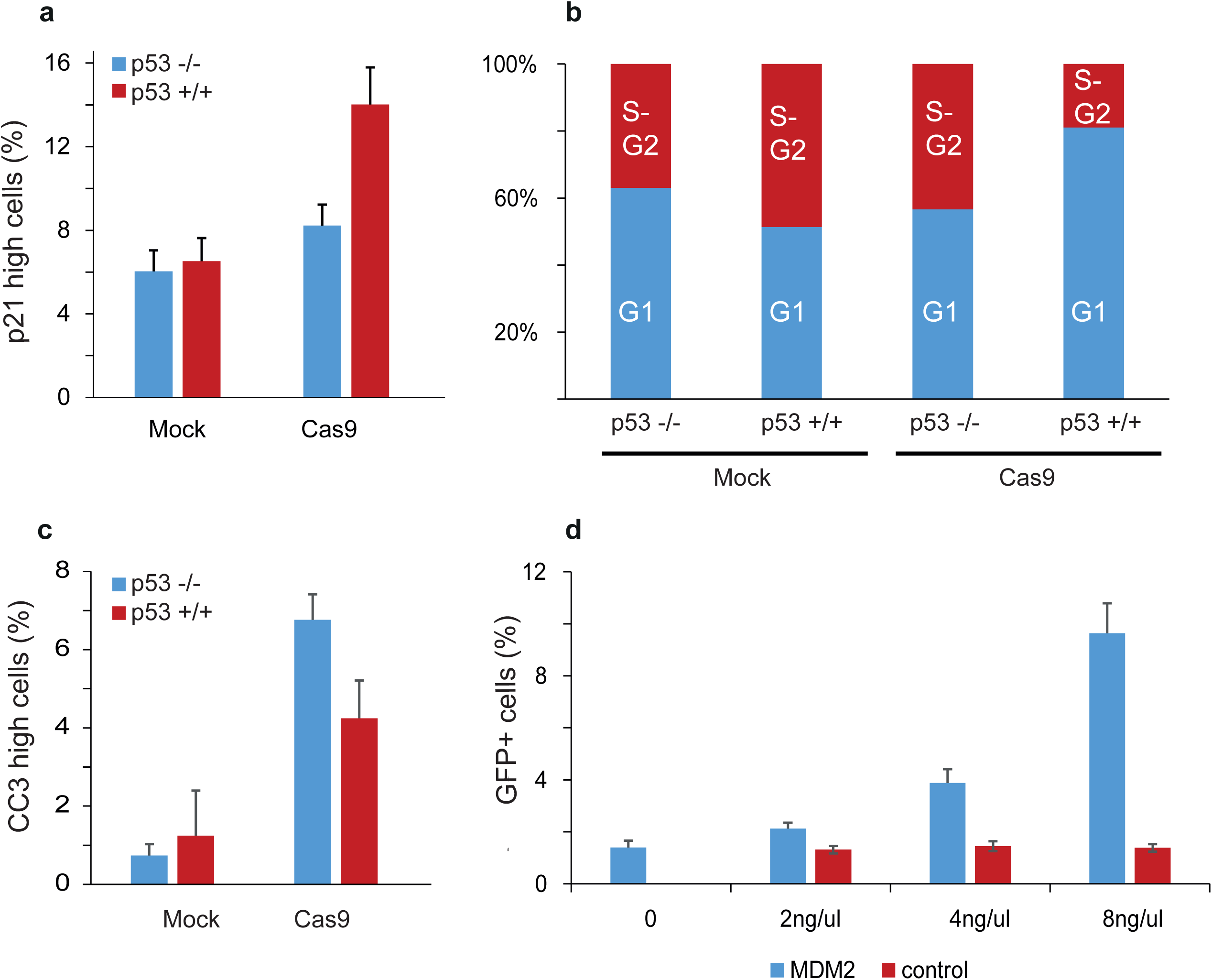
Cas9-gRNA ribonucleoprotein complex delivery triggers p53 DNA damage response. RPE1 cells with WT and null p53 alleles were transfected with Cas9 ribonucleoprotein complexes targeting the RNF1 genomic locus^13^. RNP delivery results in a significant increase of p21, the target of p53 **(a)** and G1 phase cells **(b)** in p53^+/+^ cells (n=4, p<0.05). **c)** Cleaved Caspase 3 (CC3) activity 24h after RNP transfection. The proportion of apoptotic cells is not increased in the p53^+/+^ sample (n=4). **d)** RPE1 cells expressing mutant GFP were transfected with RNPs targeting GFP along with a repair DNA template that restores the GFP function. HR efficiency was evaluated by percentage of RPE1 cells that turn GFP positive. MDM2 supplementation dramatically improves the HR rate (n=6 for each condition tested). All data was acquired by flow cytometry.

Precision genome editing requires homologous recombination, which largely occurs in S-phase^8^. Thus, the observed G1 arrest is likely to preclude efficient HR, leading to the imprecise repair by NHEJ. This would explain the high NHEJ-mediated indel count and the low efficiency of HR-dependent precision gene-editing, in particular in primary cells^1^. These results suggest that p53 inhibition should increase the frequency of homologous recombination and precision genome editing, as it would allow cell cycle progression in the presence of Cas9-induced DSBs. To test this hypothesis, we used an RPE1 p53^+/+^ reporter cell line expressing a mutationally inactivated green fluorescent protein (GFP). Co-transfecting these cells with a single-stranded DNA repair template and Cas9 RNP targeting the mutation site of the GFP restored GFP fluorescence. Co-transfecting the cells with the p53 antagonist MDM2 dramatically and dose-dependently improved the repair efficiency (Fig. 2D), indicating that triggering of cellular DNA damage response has a major role in limiting the efficiency of precision genome editing in normal human cells.

The cell growth arrest and apoptosis upon CRISPR-Cas complex delivery has also been suggested to result from type I interferon (IFN) response, which can be activated upon plasmid, protein and lentiviral transfection^2, 14, 15^. Since the Toll-like receptor, Interferon-α and Interleukin 1β signaling comprise the major branches of this pathway, we tested the effect of several established IFN inhibitors on precision genome editing using the RPE-1-GFP cell line described above (Supplementary Table S2). None of the tested inhibitors, or combinations thereof had a significant effect on efficiency of genome editing. These results suggest that type I interferon response is not a major contributor to the observed CRISPR-Cas toxicity and G1 arrest in RPE1 cells, and that its inhibition cannot be used to increase the rate of HR.

We report here that genome editing by Cas9 in p53 proficient cells results in DNA damage response, leading to preferential repair of the induced lesion by the imprecise process of NHEJ. The observed effect is dependent on p53, its downstream target p21, and RB1, which is necessary for G1 cell cycle arrest in response to p21 induction^8^. Based on failure of non-targeting gRNAs to induce growth arrest in the RPE1 screen (Fig. 1C), the effect depends on the presence of DNA double-strand breaks. The general effect of all gRNAs irrespective of their genomic targets, in turn, suggests that very few DNA lesions are sufficient for the growth arrest. Cas9 has been shown to associate with cut sites for over six hours after the double-strand break ^5^, preventing a successful repair; this is likely to amplify the effect of a single DSB, leading to a generalized DNA damage response.

Our results show that decreasing DNA damage signaling dramatically improves the efficiency of precision genome editing in normal cells. At the same time, inhibition of p53 leaves the cell transiently vulnerable to the introduction of chromosomal rearrangements and other tumorigenic mutations. It should be noted that some other reported improvements of genome editing may also sensitize cells to additional DNA damage. For example, use of adeno-associated virus (AAV) in repair DNA delivery has been reported to improve HR in human p53 proficient tumor cells and in normal hematopoietic stem cells^2, 16^. The AAV encoded gene Rep78 is a p53 binding protein that may be partially responsible for this effect^2, 18^.

Based on the findings that effective genome editing depends on lack of p53 signaling, and that the genome editing itself exerts a selective pressure that leads to a preferential growth of p53 deficient cells, we strongly advise caution in the therapeutic use of CRISPR/Cas. The temporary inhibition of p53 in normal cells will have both positive and negative effects on the tumorigenic potential of the edited cell population. On one hand, it could potentially allow escape of cells whose genome is damaged during the editing process. On the other hand, it will increase the editing efficiency, and decrease the selective advantage of pre-existing p53 deficient clones. To improve the balance, future work should focus on understanding the DNA damage response induced by Cas9. Controlling the DNA damage signaling process in such a way that allows efficient genome editing, but does not select for or facilitate the formation of potentially tumorigenic cells will be critically important for future efforts in therapeutic genome editing.

## Author contributions

E.H., B.S. and J.T. wrote the manuscript. S.B., B.S. and J.P. conducted the genome-wide knockout screens. E.H., B.S. and S.B. prepared the cell lines and performed the flow cytometry experiments. J.T. supervised the study. All authors read and approved the final manuscript.

The authors declare no conflict of interests.

## Acknowledgements

Part of this work was carried out at the High Throughput Genome Engineering Facility and the Swedish National Genomics Infrastructure funded by Science for Life Laboratory. Knut and Alice Wallenberg Foundation, Cancerfonden and Academy of Finland supported this work.

After completion of the experiments described herein, an independent manuscript was posted to BioRxiv that describes Cas9 induced and p53 dependent toxicity in human pluripotent stem cells (http://www.biorxiv.org/content/early/2017/07/26/168443). The findings described in this manuscript are consistent with our findings, and further highlight the need for caution in clinical use of currently available CRISPR/Cas9 genome editing tools.

